# Exon-Skipping Antisense Oligonucleotides for Cystic Fibrosis Therapy

**DOI:** 10.1101/2021.08.11.455936

**Authors:** Young Jin Kim, Nicole Sivetz, Jessica Layne, Dillon Voss, Lucia Yang, Qian Zhang, Adrian R. Krainer

**Affiliations:** Cold Spring Harbor Laboratory, Cold Spring Harbor, New York, USA; Department of Genetics, Stony Brook University, Stony Brook, New York, USA; Medical Scientist Training Program, Stony Brook University School of Medicine, Stony Brook, New York, USA; Graduate Program in Molecular and Cell Biology, Stony Brook University, Stony Brook, New York, USA; Stanford University, Palo Alto, CA, USA

**Keywords:** Cystic fibrosis, nonsense-mediated mRNA decay, pre-mRNA splicing, antisense oligonucleotides

## Abstract

Mutations in the cystic fibrosis transmembrane conductance regulator (*CFTR*) gene cause cystic fibrosis (CF), and the *CFTR-W1282X* nonsense mutation causes a severe form of CF. Although Trikafta and other CFTR-modulation therapies benefit most CF patients, targeted therapy for patients with the W1282X mutation is lacking. The CFTR-W1282X protein has residual activity, but is expressed at a very low level due to nonsense-mediated mRNA decay (NMD). NMD-suppression therapy and read-through therapy are actively being researched for *CFTR* nonsense mutants. NMD suppression could increase the mutant *CFTR* mRNA, and read-through therapies may increase the levels of full-length CFTR protein. However, these approaches have limitations and potential side effects: because the NMD machinery also regulates the expression of many normal mRNAs, broad inhibition of the pathway is not desirable; and read-through drugs are inefficient, partly because the mutant mRNA template is subject to NMD. To bypass these issues, we pursued an exon-skipping antisense oligonucleotide (ASO) strategy to achieve gene-specific NMD evasion. A cocktail of two splice-site-targeting ASOs induced the expression of *CFTR* mRNA without the PTC-containing exon 23 (CFTR-Δex23), which is an in-frame exon. Treatment of human bronchial epithelial cells with this cocktail of ASOs that target the splice sites flanking exon 23 results in efficient skipping of exon 23 and an increase in CFTR-Δex23 protein. The splice-switching ASO cocktail increases the CFTR-mediated chloride current in human bronchial epithelial cells. Our results set the stage for developing an allele-specific therapy for CF caused by the W1282X mutation.

## Introduction

Nonsense mutations account for about 20% of all known pathogenic genetic lesions within the coding region (1). Nonsense mutations in the cystic fibrosis transmembrane conductance regulator (*CFTR*) gene cause cystic fibrosis (CF) by impairing chloride conductance in the epithelial tissues of multiple organs, including lung, gastrointestinal tract, and reproductive organs (2). The *CFTR-W1282X* mutation is the 6^th^ most common CF-causing mutation and the 2^nd^ most common CF-causing nonsense mutation; it is found in 1.2 % of CF patients worldwide, and in up to 40% of Israeli CF patients, and causes a severe form of CF if homozygous or combined with another CF-causing allele (3, 4). Nonsense mutations, including *CFTR-W1282X* mutation, lead to reduction of functional proteins due to premature termination of translation and degradation of the mRNA due to nonsense-mediated mRNA decay (NMD) (5).

There have been significant strides in CF therapeutics development, with the most recent FDA-approved treatment, Trikafta—a combination of CFTR correctors (elexacaftor and tezacaftor) and a potentiator (ivacaftor)—showing unprecedented improvements in the pulmonary exacerbations, survival, and quality of life for CF patients with at least one *F508del* mutation (6). Although almost 90% of CF patients may benefit from TRIKAFTA, no treatments currently exist for CF patients not carrying the *F508del* mutation. Gene therapy is another actively explored strategy for CF caused by nonsense mutations, and promising results were shown in various model systems; however, various challenges, including inefficient delivery and gene expression have to be overcome before they can be used as therapy (7). Therefore, there is still a significant unmet need for CF treatments that can simultaneously address different classes of CFTR protein deficiencies, particularly the *CFTR-W1282X* mutation (8).

One potential therapeutic modality for *CFTR* nonsense mutations is read-through compounds, which increase the level of full-length protein by reducing the fidelity of the ribosome at the premature termination codon (PTC) (9). Ataluren is a non-aminoglycoside RTC with a favorable safety profile that was investigated in the context of several genetic diseases. Although it showed a therapeutic effect in Duchenne muscular dystrophy caused by nonsense mutations (9), it was not able to improve forced expiratory volume (FEV) in clinical trials in CF patients with various *CFTR* nonsense mutations (10). Gentamicin is an aminoglycoside RTC that can increase full-length CFTR protein *in vitro,* whose clinical efficacy in CF is limited by NMD (11). Recently, a new class of RTC that induces readthrough by depleting the translation release factor eRF1 showed efficacy *in vitro* (12). Despite active research, a clinically viable read-through approach for CF caused by nonsense mutation does not yet exist.

The truncated CFTR-W1282X protein is weakly functional, and *CFTR-W128X* mRNA expression is very low, due to the PTC in exon 23 that targets the *CFTR-W128X* transcript for NMD (4, 13, 14). Because the activity of CFTR-W1282X protein can be enhanced by CFTR modulators (4, 13, 14), combining NMD-inhibition strategies with CFTR modulators could potentially benefit patients. In one such strategy, an inhibitor of the SMG1 kinase that efficiently suppresses NMD in human bronchial epithelial cells increases expression and function of CFTR in cells harboring the *W1282X* mutation (15). Another strategy involves antisense oligonucleotides, which are short, synthetic, single-stranded DNA, RNA, or structural mimics of DNA/RNA that bind to complementary RNA to alter its functions (16). ASOs with uniformly modified nucleotides, such as 2’-O-methoxyethylribose-phosphorothioate (PS-MOE-ASOs), phosphorodiamidate morpholino oligomer (PMO), or 2’-O-Methyl (2’-OMe), can sterically block the binding of proteins, nucleic acids, or other factors that are important for RNA metabolism and processing (16). On the other hand, chemically modified nucleotides in ‘gapmer’ ASOs are spaced out by natural DNA nucleotides, in order to induce RNase-H-mediated degradation of their target RNA (16). Antisense-mediated knockdown of *SMG1* inhibits NMD in human bronchial epithelial cells, and improves expression and function of CFTR in cells harboring the *W1282X* mutation (15). However, general inhibition of NMD may not be the optimal strategy, because the NMD machinery regulates expression of a subset of normal and physiologically functional mRNAs (17). In addition, some NMD factors are essential for embryonic and post-embryonic vertebrate development, and mutations in them cause developmental disorders in humans (17). Therefore, a targeted approach for restoring functional *CFTR* mRNA seems a more desirable therapeutic strategy.

According to the ‘55-nucleotide rule’, mRNA with an exon-exon junction >55 nucleotides (nt) downstream of a stop codon is degraded by NMD, due to the binding of exon-junction complexes (EJC) near each of the downstream exon-exon junctions (5). Attempts have been made to target the multiple exon-exon junctions located downstream of the PTC on the *CFTR-W1282X* mRNA to inhibit NMD. Deleting the genic region downstream of the W1282X mutation using CRISPR-Cas9 genome editing prevents NMD of *CFTR-W1282X* mRNA and increases the truncated protein levels in human airway cells (18). PS-MOE ASOs designed to prevent EJC binding downstream of a PTC can attenuate NMD in a gene-specific manner (19). Recently, we showed that a cocktail of three ASOs designed to prevent EJC binding downstream of the W1282X mutation specifically increases the expression of *CFTR*-*W1282X* mRNA and CFTR protein function in human bronchial epithelial cells (20).

An alternative strategy we describe here is to upregulate a partially functional isoform of *CFTR* mRNA using ASOs that can modulate mRNA splicing. In the recent years, ASOs drugs have been shown to be well-suited for treatment of various diseases (21). The clinical potential of ASOs for hereditary diseases has been demonstrated by recent FDA/EMA regulatory approvals. Several PMO ASOs (eteplirsen, golodirsen, viltolarsen, and casimersen) have been approved to treat different causal mutations in Duchenne muscular dystrophy; they induce skipping of target exons in the mutant *DMD* pre-mRNA and promote the production of partially functional dystrophin protein (22). The FDA approval of nusinersen (Spinraza) in 2016 to treat spinal muscular atrophy and the rapid development of milasen, a personalized therapy for a single patient with neuronal ceroid lipofuscinosis 7 (CLN7, a form of Batten’s disease), further highlight the clinical potential of splice-switching ASOs (23, 24). Although there is no FDA-approved ASO therapy for CF at present, splice-switching ASOs designed to correct *CFTR* defective splicing have shown efficacy in patient-derived airway cells and nasal epithelial cells (25, 26).

If *CFTR* exon 23 with the W1282X mutation is skipped using splicing-switching ASOs, the resulting *CFTR-*Δ*ex23* mRNA should not be targeted by NMD, as exon 23 is in-frame. The intrinsic activity of CFTR-Δex23 has not been reported, but it is expected to be less active than wild-type CFTR, based on the deletion’s location in the second nucleotide-binding domain (NBD2). On the other hand, this protein is expected to retain partial activity, by analogy to the truncated CFTR-W1282X protein. CF-causing exon 23 splice-site mutations (4005+1G>A and +2T>C) have been identified (3); however, the intrinsic activity of CFTR-Δex23 cannot be inferred from the phenotypes of these mutations, because whether they cause exon skipping, cryptic splice-site activation, and/or intron retention is unknown (27).

As mentioned above, CFTR modulators can increase the activity of a wide variety of mutant CFTR proteins (28), so CFTR-Δex23 protein’s chloride conductance may also be enhanced by CFTR modulators. Moreover, as low as 10% of normal CFTR function provides a significant therapeutic benefit for CF patients (29). Thus, increasing the expression of mutant CFTR protein with residual activity could be beneficial for CF patients. In this report, we show that CFTR-Δex23 protein retains partial activity. We also describe uniformly modified PS-MOE ASOs that target the splice sites flanking exon 23 and induce efficient exon 23 skipping, with the resulting CFTR-Δex23 protein increasing CFTR-mediated chloride conductance in human bronchial epithelial cells. This approach provides the basis for clinical development of antisense-directed exon skipping as a therapeutic strategy for CF caused by the W1282X mutation.

## Results

### Intrinsic activity of CFTR-Δex23

To assess the functionality of CFTR protein lacking the internal peptide (amino acids 1240-1291) encoded by exon 23 (CFTR-Δ ex23), we generated 16HBE-W1282X cells with doxycycline (dox)-inducible overexpression of wild-type CFTR-WT or CFTR-Δex23 (dubbed 16HBEge-GFP-P2A-WT or 16HBEge-GFP-P2A-Δex23), using a lentiviral vector that allows co-expression of Turbo GFP using a 2A self-cleaving peptide (Fig. 1A). Dose-dependent induction of the recombinant gene expression by dox was confirmed by fluorescence microscopy (Fig. 1B-D, Fig. S1A). Since the Turbo GFP and CFTR proteins are translated initially as the same polypeptide chain, GFP levels can be used for controlling for expression from the integrated plasmid (30). Treating the 16HBEge-GFP-P2A-WT and 16HBEge-GFP-P2A-Δex23 cells with 2 μg/mL doxycycline resulted in similar levels of GFP signal, indicating that the expression of recombinant CFTR proteins is induced at similar levels (Fig. S1B).

**Figure 1.**
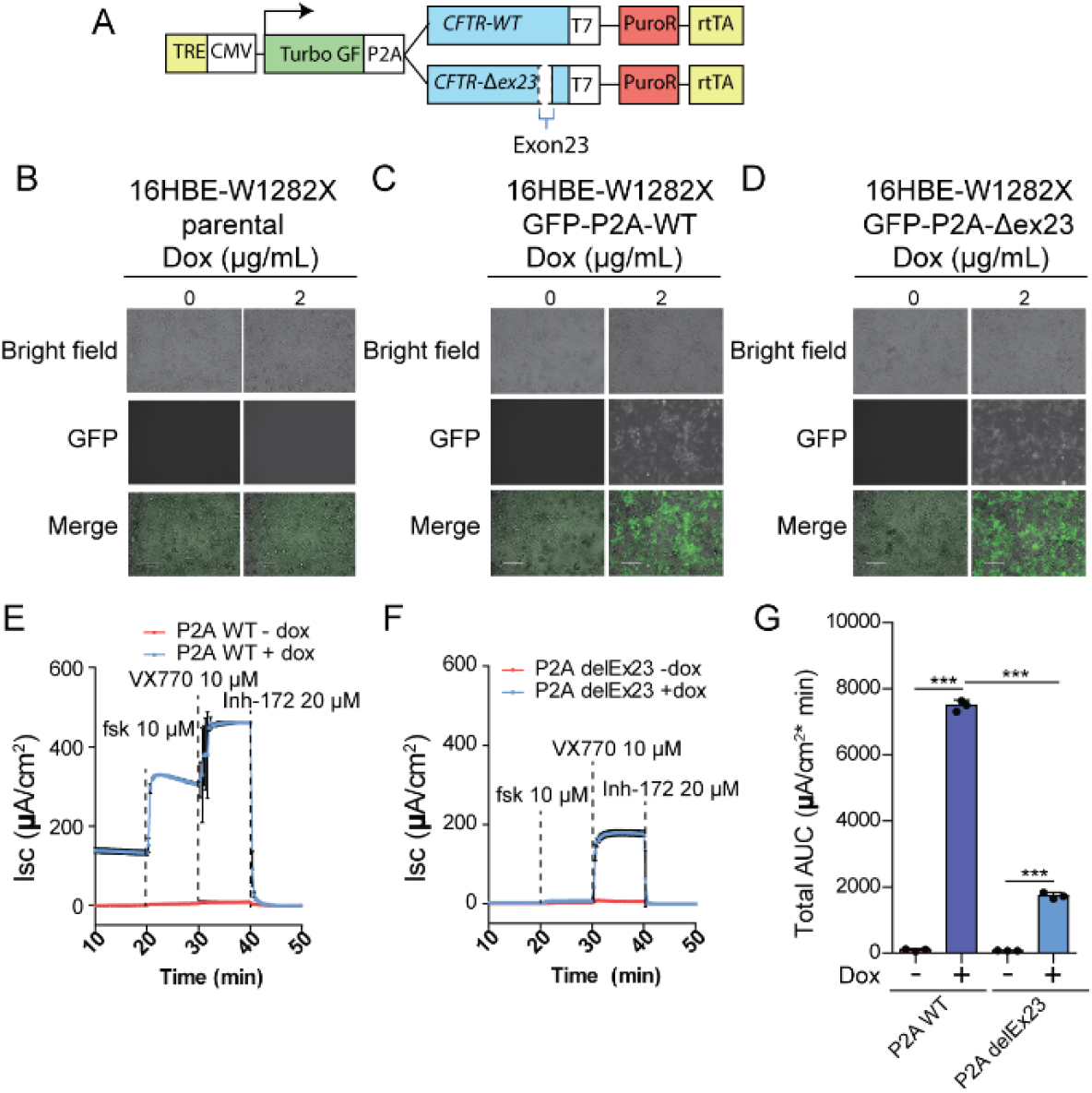
Intrinsic activity of CFTR-Δex23 protein. **A.** Schematic of recombinant CFTR expression system. **B-D.** Fluorescence microscopy images of (B) 16HBE-W1282X cells, (C) 16HBEge-GFP-P2A-WT, and (D) 16HBEge-GFP-P2A-Δex23 cells treated with doxycycline for 48 hours. The white bars are 120-μm rulers. **E-F.** Average traces from Ussing-chamber assays of (E) 16HBEge-GFP-P2A-WT and (F) 16HBEge-GFP-P2A-Δex23 cells treated with doxycycline and TRIKAFTA. The traces shown are the averages of three replicates. Black error bars on each point show standard deviation. **G.** The total area under the curve of (F) between minutes 20-40 (n=3 independent treatments. n.s. P>0.05, *P<0.05, **P<0.01, ***P<0.001, Student’s t-test). Error bars show standard deviations. TRE = tetracycline-response-element. CMV = CMV promoter. P2A = self-cleaving 2A peptide. PuroR = puromycin-resistance gene. rtTA = reverse tetracycline transactivator. Dox = doxycycline. Fsk: forskolin.

Confocal immunofluorescence microscopy showed that the T7-tagged recombinant CFTR-WT and CFTR-Δex23 proteins are localized to the cytoplasm and plasma membrane (Fig. S1C-E). We compared the chloride currents with CFTR-WT or CFTR-Δex23 overexpression by Ussing chamber assays. 16HBEge-GFP-P2A-WT or 16HBEge-GFP-P2A-Δex23 cells treated with dox and TRIKAFTA (VX-445, VX-661, and VX-770) led to significant increases in chloride current compared to TRIKAFTA only treatment, as quantified by the total area under the curve (Fig. 1E-G). The total AUC of 16HBEge-GFP-P2A-Δex23 after VX-770 treatment was about 53% and 38% of the total AUC before and after VX-770 treatment in 16HBEge-GFP-P2A-WT cells, respectively (Fig. S2). The background CFTR activity in 16HBE-W1282X cells due to endogenous CFTR-W1282X expression is negligible, as the chloride currents of the no-doxycycline-treatment 16HBEge-GFP-P2A-WT (red traces in Fig. 1E) and 16HBEge-GFP-P2A-Δex23 (red traces in Fig. 1F) cells are substantially lower than those of the doxycycline-treated cells (blue traces in Fig. 1E-F). We conclude that the CFTR-Δex23 protein is partially active.

### Design and testing of candidate ASOs

ASOs targeting ESEs or splice sites can induce skipping of alternative or constitutive exons (31, 32). We designed two PS-MOE ASOs targeting the splice sites flanking exon 23 (i22-3ss and i23-5ss). We used splicing-factor binding-site predictions by SFmap (Fig. S2A) and ESEfinder (Fig. S2B) to design three additional ASOs targeting potential ESEs (33, 34). We tested these ASOs in 16HBE-W1282X cells, in which the endogenous *CFTR-W1282X* mRNA is degraded by NMD (Fig. 2A-B). 16HBE-W1282X cells transfected with the splice-site-targeting ASOs increased exon 23 skipping significantly, compared to the scramble negative control ASO. The identity of the exon-23-skipped bands detected by RT-PCR was confirmed by Sanger sequencing (Fig. S2C). Though we could detect exon 23 skipping in the presence of ESE-1 and ESE-2 ASOs, the decrease in exon-23-included mRNA was not statistically significant for any of the three ASOs designed to target potential ESEs (Fig. 2C-D). These results suggest that the target sites for ASOs ESE-1/2/3 are not strong binding sites for splicing activators, or they are not individually essential for exon recognition. Thus, we selected the ASOs that target the 3’ss and 5’ss as the lead ASOs.

**Figure 2.**
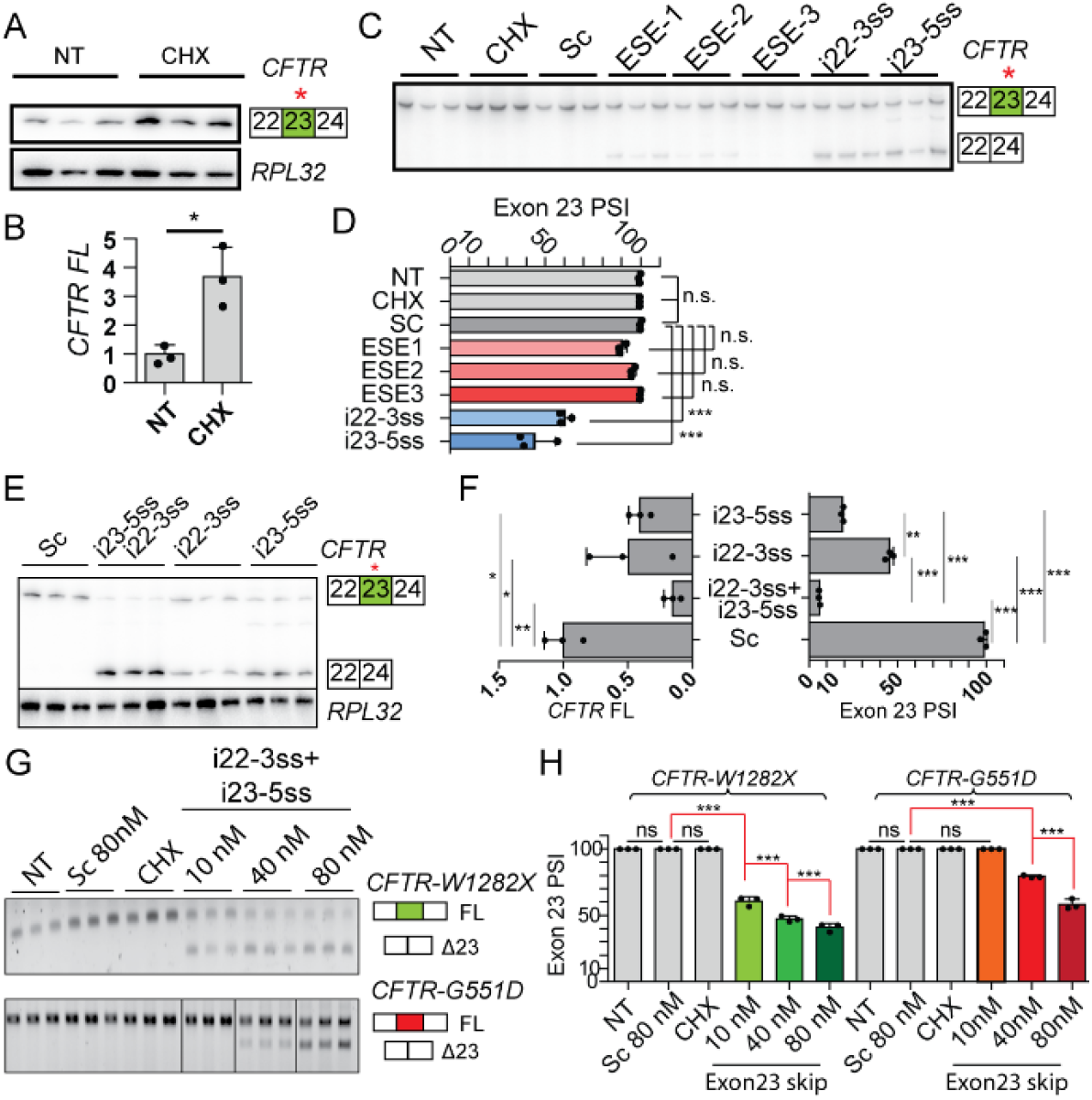
Designing ASOs that induce exon 23 skipping. **A.** RT-PCR of *CFTR-W1282X* mRNA in 16HBE-W1282X cells with NT and CHX treatment using *RPL32* as internal reference. **B.** Quantification of (A). **C.** RT-PCR of *CFTR* mRNA in 16HBE-W1282X transfected with the ASOs complementary to the splice sites flanking *CFTR* exon 23 at a nominal concentration of 80 nM. **D.** Quantification of exon 23 inclusion based on (D). **E.** RT-PCR comparing the exon 23 PSI between the transfection of individual splice-site-targeting ASOs (80 μM) and the ASO cocktail (40 μM each ASO). **F.** Exon 23 PSI and full-length *CFTR-W1282X* mRNA based on (E). **G.** Non-radioactive RT-PCR dose-dependent exon 23 skipping in 16HBE-W1282X and 16HBE-G551D cells treated with splice-site ASO cocktail. All samples were run on the same gel, but some lanes were rearranged for presentation. **H.** Exon 23 PSI based on (G). Mean PSI and standard deviation in parentheses of exon 23 are shown below. *RPL32* as an internal reference for panels B and F. For all treatments, n=3 independent transfections. For all statistical tests, n.s. P>0.05, *P<0.05, **P<0.01, ***P<0.001. Panel B: Student’s t-test. Right half of panel F and panel H: one-way ANOVA with Tukey’s post-test. Left half of panel F: one-way ANOVA with Dunnett’s post-test. Error bars are standard deviations. PSI = percent-spliced-in. Sc = scramble ASO. NT = No-treatment control. CHX = cycloheximide treatment.

### Combination of splice-site ASOs induces efficient exon 23 skipping

To test whether targeting the 3’ss and 5’ss simultaneously promotes efficient exon skipping, we transfected 16HBE-W1282X cells with ASOs i22-3ss and i23-5ss individually or as a cocktail. The splice-site ASO cocktail resulted in almost complete skipping of exon 23, compared to the partial skipping elicited by the individual ASOs and significantly reduced the level of full-length *CFTR* mRNA, compared to the scramble ASO (Fig. 2E-F). The splice-site ASO cocktail increased exon 23 skipping of in 16HBE-W1282X cells in a dose-dependent manner (Fig. 2G-H). As the W1282X (TGG -> TGA) mutation disrupts potential binding sites for SRSF1 and SRSF2, according to ESEfinder (Fig. S2A-B), we tested if the ASO cocktail promotes exon 23 skipping in *CFTR-G551D* mRNA, which does not have a mutation on exon 23. The lead ASO cocktail also promoted dose-dependent exon 23 skipping in 16HBE-G551D cells, but the extent of exon 23 skipping was lower in these cells (Fig. 2H). Treating the 16HBE-W1282X cells with 10 μM ASO cocktail by free uptake also induced efficient exon skipping (Fig. S3).

However, calculating the proportion of exon 23 inclusion without taking mRNA degradation into account can overestimate the splice-site ASO cocktail’s efficacy. Because the full-length *CFTR-W1282X* mRNA is an NMD target, we compared the extent of exon 23 inclusion between 16HBE-W1282X cells treated with the ASO cocktail only versus the ASO cocktail plus cycloheximide (to inhibit NMD). The combined treatment significantly increased the exon 23 PSI, compared to the ASO cocktail treatment alone (Fig. 3A-B). Compared to the controls, the full-length *CFTR-W1282X* mRNA levels were significantly lower for the ASO cocktail alone or in combination with cycloheximide (Fig. 3C). RT-qPCR using a primer pair complementary to *CFTR* exons 26 and 27 to amplify both full-length and exon 23-skipped isoforms shows that the ASO cocktail increased the total *CFTR* mRNA in 16HBE-W1282X cells (Fig. 3D). Combining the ASO cocktail with cycloheximide also increased the total *CFTR* mRNA (Fig. 3D). These results show that the splice-site ASO cocktail efficiently induces *CFTR* exon 23 skipping, bypassing NMD and resulting in an increase in the total *CFTR* mRNA levels (Fig. 3E).

**Figure 3.**
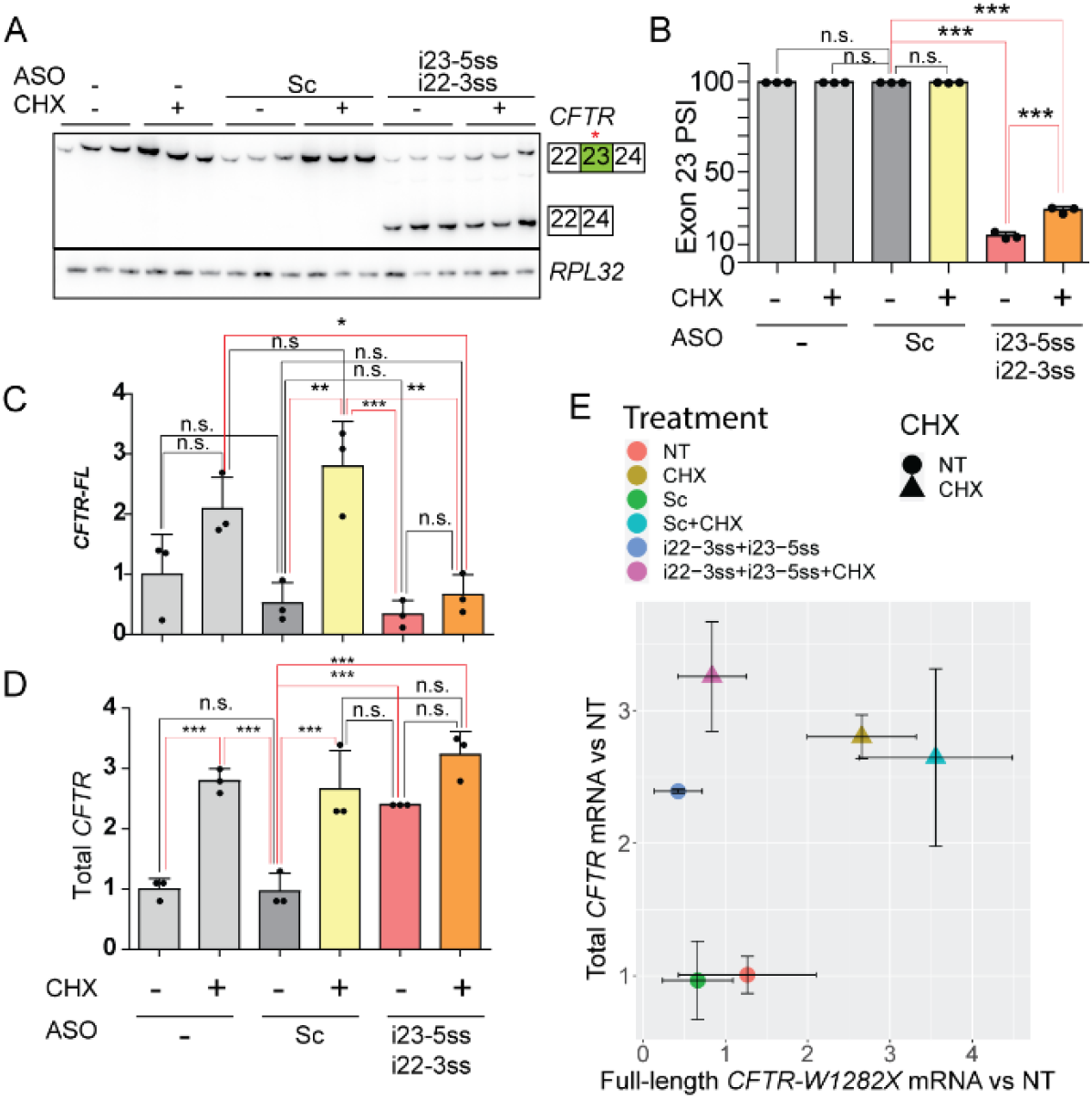
RT-PCR and RT-qPCR analysis of total *CFTR* mRNA and exon 23 splicing after treatment with the splice-site ASO cocktail alone or in combination with NMD inhibition. **A.** NMD inhibition was combined with ASO treatments. **B-C.** Quantification of (B) exon 23 PSI and (C) full-length *CFTR-W1282X* mRNA based on (A). **D.** Total *CFTR* mRNA levels (full-length *CFTR-W1282X* and *CFTR-*Δ*ex23* mRNA) were measured by RT-qPCR. **E.** The full-length *CFTR-W1282X* mRNA levels (from C) was plotted against the total *CFTR* mRNA levels (from D). Circle dots show no CHX treatment, and triangle dots show CHX treatment. *RPL32* mRNA levels were used as an internal reference for (C) and (D). For all treatments and statistical tests, n=3 independent transfections, n.s. P<0.05, ***P<0.001, one-way ANOVA with Tukey’s post-test. Error bars are standard deviations. Abbreviations are as in Fig. 2.

### The splice-site ASO cocktail increases CFTR-Δex23 protein and CFTR activity

As the splice site ASO cocktail effectively promoted *CFTR* exon 23 skipping, we next analyzed the expression and function of the resulting CFTR-Δex23 protein. Three different glycosylation states of CFTR-WT can be seen on a Western blot: non-glycosylated A-band (127 kDa), core glycosylated B-band (131 kDa), and fully mature glycosylated C-band (160-170 kDa) (35). The predicted size of fully glycosylated CFTR-Δex23 C-band is about 160 kDa. Transfection or free-uptake delivery of the splice-site ASO cocktail into 16HBE-W1282X cells increased the level of CFTR protein (Fig. 4A-D). Whereas the DLD1 cell-lysate sample shows two CFTR-WT bands (C-band and B-band), the protein samples from the 16HBE-W1282X cells treated with the splice-site ASO cocktail show one predominant band that is presumably a CFTR-Δex23 B band, based on its size (Fig. 4B). An antibody that can specifically detect the part of CFTR encoded by exon 23 does not exist, to our knowledge. However, as shown above, the splice-site ASO cocktail decreased the full-length *CFTR-W1282X* mRNA, so the increased CFTR protein is most likely the CFTR-Δex23 protein.

**Figure 4.**
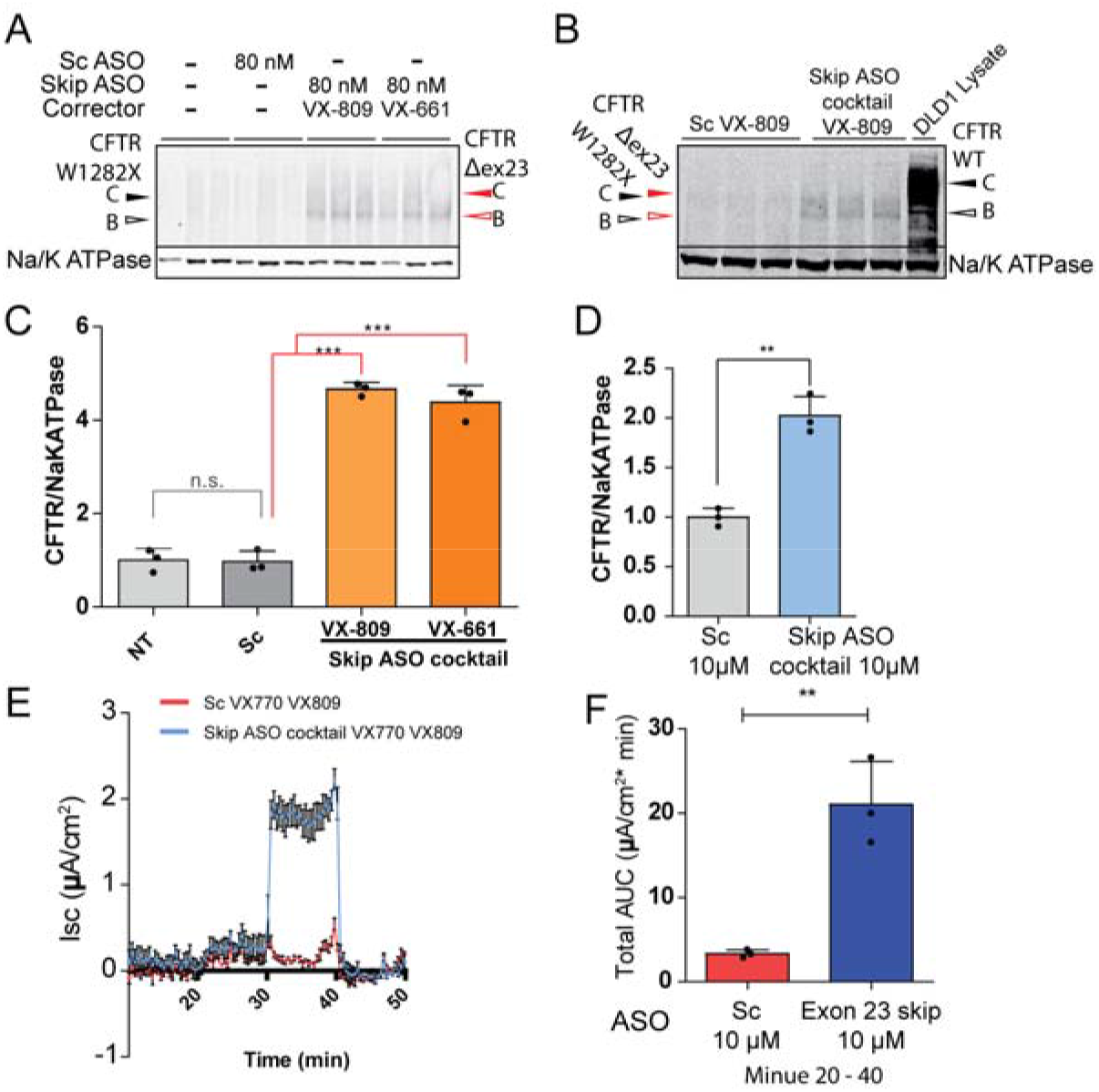
Combining the splice-site ASO cocktail and CFTR modulators increases CFTR-Δex23 protein levels and CFTR activity. **A-B.** Western blot of 16HBE-W1282X cells treated with the splice-site ASO cocktail by (A) transfection or (B) free uptake. Na/K ATPase was used as an internal reference. The open and filled arrowheads represent core-glycosylated and complex-glycosylated CFTR, respectively. **C-D.** Quantification of (A) and (B). **E.** The data points represent the mean I_sc_ tracings of three independent 16HBE-W1282X cells treated with Sc ASO (red) or the splice-site ASO cocktail (blue). Both ASO treatments were combined with 10 μM VX-770 and 5 μM VX-809 treatments. **F.** The total area under the curve of (E). For all treatments, n=3 independent treatments. For all statistical tests, n.s. P>0.05, **P<0.01, and ***P<0.001. Panel C: one-way ANOVA with Tukey’s post-test. For panel D and F: Student’s unpaired t-test, versus Scramble ASO. Error bars are standard deviations. Abbreviations are as in Fig. 2.

We measured CFTR function in 16HBE-W1282X cells treated with the splice-site ASO cocktail by Ussing chamber assay. The 16HBE-W1282X cells treated with the splice-site ASO cocktail by free uptake in combination with VX-809 and VX-770 showed increased CFTR-mediated chloride current, compared to the cells treated with scramble 18mer ASO (Fig. 4E-F). The increase in CFTR activity was similar to what we previously observed by treatment with an EJC-targeting triple-ASO cocktail, plus VX-809 and VX-770 (39).

## Discussion

Although major advances have been made for small-molecule therapies for CF that predominantly benefit patients with at least one *CFTR-F508del* allele, therapeutic options are lacking for patients harboring nonsense mutations in *CFTR* (6, 36). CFTR nonsense mutations, including *CFTR-W1282X*, lead to low levels of functional protein, due to NMD and premature termination of translation. Overcoming NMD without disrupting the physiological gene-regulatory role of this pathway is a key objective (17). Although allele-specific NMD-inhibition strategies were recently developed, using genome editing with CRISPR-Cas9 or ASOs that block EJC deposition (18, 20), these preclinical studies have yet to be translated to the bedside. Thus, there is an unmet need for therapies for CF caused by nonsense mutations. As low as 10% of normal CFTR function would provide a significant therapeutic benefit for CF patients who have a near complete loss of CFTR function, as is the case for the W1282X mutation (29). Thus, increasing the expression of mutant CFTR protein with residual activity using splice-switching ASOs could be beneficial for CF patients, as long as CFTR-Δex23 protein retains some function.

Compared to other gene-specific NMD inhibition approaches using CRISPR-Cas9 genome editing or ASOs that inhibits EJC binding (GAIN) (15, 37), inducing expression of CFTR-Δex23 protein may have several advantages. CFTR-Δex23 protein retains C-terminal-domain residues (^1478^TRL^1480^) that are important for post-translational processing and CFTR gating, which are missing in the C-terminal-truncated CFTR-W1282X protein (38–40). ATP binding and dimerization of NBD1 and NBD2 are critical for CFTR gating (41). Regions important for ATP binding, namely the Walker B motif (residues 1366 to 1372) and the NBD2 signature sequence (residues 1346 to 1350) are present in CFTR-Δex23 protein (which lacks residues 1240-1291) but are missing in the truncated W1282X mutant protein.

Our recently described EJC-targeting ASO approach can effectively inhibit NMD, but requires a cocktail of three ASOs that block individual EJC binding sites downstream of the W1282X codon (19). For lead compound optimization, the EJC-targeting approach requires optimization of length, chemical modifications, and dose, as well as analysis of toxicity, for each of the ASOs in the cocktail. On the other hand the exon-skipping ASO strategy can potentially be achieved using a single ASO that targets an ESE or intronic splicing enhancer (ISE) element. Our limited attempt to identify exon-skipping ASOs targeting putative ESEs in exon 23 was not successful, but we expect that effective exon-skipping ASOs may be identified through a systematic ASO screen of exonic and intronic regions, as described previously for other pre-mRNAs (31, 32).

Based on the measurement of CFTR-Δex23 activity, while controlling for the plasmid expression levels, we estimate that CFTR-Δex23 protein has residual activity as high as 54% of WT, when combined with TRIKAFTA. We show that uniform PS-MOE ASOs targeting the pre-mRNA 3’ss or 5’ss flanking *CFTR* exon 23 induce exon 23 skipping. Targeting both splice sites simultaneously with an ASO cocktail induced more efficient exon 23 skipping than targeting each individual splice site with a single ASO. Measuring the exon 23 PSI after cycloheximide treatment to inhibit NMD showed that the lead ASO cocktail caused almost complete exon 23 skipping. The *CFTR-*Δ*ex23* mRNA wasre not sensitive to NMD inhibition by cycloheximide, and it was translated into CFTR-Δex23 protein that retained partial function in human bronchial epithelial cells. Compared to the published activity of WT CFTR in 16HBE17o- cells (37), the 16HBE-W1282X cells treated with the splice-switching lead ASO cocktail combined with a CFTR corrector showed < 10% of the WT activity. We anticipated higher CFTR activity in these cells, given our results with the recombinant CFTR-Δex23 expression experiments, but several factors may be at play. Compared to the delivery of ASO by transfection, free uptake of naked ASOs is less efficient. ASO free uptake occurs mainly through receptor-mediated endocytosis, and only a small portion of the ASOs escapes from endosomes and can reach the nucleus to engage its RNA target (42). In addition to optimizing delivery, further optimization of the ASOs in terms of chemistry and dosing could be helpful for improving the rescue of CFTR function by the ASO cocktail.

ESEfinder and SFmap predictions are not sufficiently robust to obviate systematic ASO screening, but they can help to establish the mechanism of action of ASOs. We observed that transfecting the lead ASO cocktail at 10 nM caused more exon skipping in 16HBE-W1282X cells than in 16HBE-G551D cells (Fig. 2H). Although the extent of exon skipping in 16HBE-W1282X may have been overestimated, as we did not use any NMD inhibitors for this experiment, this alone does not explain the lack of exon-skipped isoform in 16HBE-G551D cells at the low ASO concentration. ESEfinder analysis predicts that the W1282X mutation may lead to the loss of SRSF1 and SRSF2 binding sites (Fig. S2B), which in turn suggests a sensitized background with a reduced threshold to undergo exon skipping in response to ASOs targeting the splice sites.

*CFTR* nonsense mutations account for about 7% of all CF-causing mutations listed in the CFTR2 database (3). More common nonsense mutations, including *G542X* and *W1282X*, are known to trigger NMD (20), and less common nonsense mutations— except for *S1455X* and *Q1476X*, which are located on the terminal exon—likely trigger NMD, according to the ‘55-nt rule’. Among these, the nonsense mutations that occur on in-frame exons may benefit from a similar ASO-mediated exon skipping strategy to avoid NMD and increase partially active CFTR. However, this would depend on whether the skipped exon encodes a peptide that is dispensable for CFTR folding, localization, and activity.

The next stage in the development of the splice-switching ASOs as a potential therapy will require optimization of the delivery to the target tissues in mice. Organs affected by CF include the airway, pancreas, colon, small intestine, testes, and skin. ASO delivery to these tissues by parenteral administration is not effective, with the exception of intestinal tissues (43). PS-MOE ASOs can be effectively delivered parenterally to the liver (hepatocyte and Kupffer cells), adipocytes, kidney, bone, bone marrow, intestines, and macrophages (43).

Systemic delivery is not optimal for delivery of ASOs to the main target tissue for our approach, which is the airway, specifically ciliated airway epithelial cells in the upper airway. In mice, IV bolus injection of 5 mg/kg ASO yields a concentration in the lung of only 2 μg/g of tissue (43). To test whether our lead ASO cocktail increases CFTR expression and function in the airway epithelial cells of mice, two modes of ASO delivery can be used: intratracheal instillation and aerosolized ASO inhalation. In intratracheal instillation, a tested compound is instilled directly in the airway using a bulb-tipped gavage needle or aerosolizing microsprayer, with the assistance of an operating otoscope (44). An advantage of intratracheal instillation is the relatively easier dose control than aerosolized drug delivery. For example, gapmer ASOs targeting *Scnn1a* (encoding mouse ENaC protein) delivered by intratracheal instillation effectively knock down *Scnn1a* in the airway epithelia of a CF-like *Nedd4L* KO mouse model (45).

Aerosolized ASO is a more clinically relevant mode of delivery. Aerosolized ASOs generated with a commercial nebulizer can be effectively delivered to the lung tissue (46). The bioavailability of an aerosolized MOE ASO was 1260 %, compared to that of IV administration in mice, and reported ASO half-lives in the lung are 4 days and >7 days in mice and monkeys, respectively (43). Thus, infrequent aerosolized dosing may maintain a sufficient tissue concentration of ASOs. In *Nedd4L* KO mice, 0.1-0.3 mg/kg of gapmer ASO targeting *Scnn1a* mRNA, which resulted in 10 μg/g lung tissue concentration, reduced expression by at least 70% (47). In these mice, up to 2 mg/kg of the *Scnn1a* ASO administered by nebulizer were well tolerated, with minimal systemic exposure. In general, the PS or PS-MOE ASOs have a very good toxicology profile in mice (46). In a phase-1 clinical trial, the PS-MOE gapmer AIR645 (Altair/Ionis), which targets IL-4/IL-13, was safely administered aerosolized at 30 mg/kg, with some adverse events (48). As the proinflammatory responses are sequence- and chemistry-specific, careful selection of ASOs will be necessary to minimize side effects.

There are currently three inhaled ASOs, developed by three companies, in active clinical trials. These developments demonstrate that aerosolization is potentially a viable delivery method to target the CF airway with our lead ASO cocktail. IONIS-ENaCRx is a cET PS gapmer ASO targeting *SCNNA1*, which completed a phase-1/2a clinical trial. TPI ASM8, from Topigen, is a combination of two PS ASOs, TOP004 and TOP005, which simultaneously target CCR-3, IL-3, IL-5, and GMCSF, which are relevant targets for asthma (49). Phase-1 and phase-2 clinical trials of TPI ASM8 were completed in 2013. These ASOs all showed in vivo activity without severe side effects (43). QR-010 is a uniformly 2’-OMe PS modified ASO developed by ProQR. QR-010 converts *CFTR-F508del* mRNA into WT *CFTR* mRNA by inserting the three missing bases in the mutant mRNA through an unknown mechanism (50). This drug is currently in a phase-2 clinical trial. The results of these studies will provide useful information about nucleic-acid therapeutics for CF and other lung diseases.

In summary, we have shown that ASO treatment directed to the 3’ss and 5’ss flanking the constitutive exon 23 of *CFTR-W1282X* pre-mRNA increases CFTR protein missing residues 1240-1291, and consequently increases CFTR activity. Further optimization of the ASOs for delivery, dosage, chemical modifications, and sequence may increase CFTR activity to a level predicted to be clinically beneficial in CF patients. This conclusion is supported by our results comparing the relative activities of CFTR-WT and CFTR-Δex23 proteins in 16HBE-W1282X cells, which showed that the CFTR-Δex23 protein in combination with TRIKAFTA has substantial activity. Our results thus provide a new avenue for developing a therapeutic strategy based on splice-switching ASO, in the era of CFTR modulator therapy.

## Materials and Methods

### CFTR expression plasmid

The plasmid pcDNA5.FRT-wtCFTR containing the coding sequence for the wild-type *CFTR* gene was kindly provided by Cystic Fibrosis Foundation’s CFFT Lab. The plasmid contains the *CFTR* gene coding sequence in the pCDNA5 FRT/TO plasmid backbone (Life Technologies, Carlsbad, CA). The *CFTR* coding sequence with a C-terminal T7 tag from pcDNA5.FRT-wtCFTR was cloned into the multiple cloning site of the pCW57-GFP-2A-MCS plasmid, resulting in the plasmid pGFP-P2A-CFTR-WT-T7. The pCW57-GFP-2A-MCS plasmid was a gift from Adam Karpf (Addgene plasmid # 71783; http://n2t.net/addgene:71783; RRID:Addgene_71783). pGFP-P2A-CFTR-WT-T7 was mutagenized to delete the coding sequence in *CFTR* exon 23, resulting in pGFP-P2A-CFTR-Δex23-T7. Plasmid constructions were conducted by GenScript (Piscataway, NJ, USA). The plasmids pGFP-P2A-CFTR-WT-T7 and pGFP-P2A-CFTR-Δex23-T7 encode recombinant GFP-P2A-CFTR proteins, which are Turbo-GFP and T7-tagged CFTR polypeptides linked by a self-cleaving 2A peptide (30).

### Tissue culture

Gene-editing of 16HBE14o- parental cells at the endogenous *CFTR* locus yielded 16HBEge cell lines CFF-16HBEge CFTR W1282X and G551D, homozygous for the *CFTR-W1282X or G551D* mutation, respectively. These cells were kindly provided by the Cystic Fibrosis Foundation’s CFFT Lab (37). Elsewhere in the text, these cells are referred to as 16HBE-W1282X and 16HBE-G551D, respectively. 16HBE-W1282X cells stably expressing pGFP-P2A-CFTR-WT-T7 or pGFP-P2A-CFTR-Δex23-T7 are dubbed 16HBEge-GFP-P2A-WT or 16HBEge-GFP-P2A-Δex23, respectively. 16HBEge cells were cultured in minimal essential medium (Thermofisher, 11095072) with 10% fetal bovine serum (Thermofisher, 26400044). DLD1 cells were cultured in DMEM with 10% FBS. The human cell line HEK293T was obtained from the American Type Culture Collection (ATCC) and cultured according to the provider’s instructions. All cells were incubated at 37 °C and 5% CO_2_. NMD inhibition by cycloheximide was performed by treating the cells for 1 hr with cycloheximide (Sigma, 100 μg/mL).

### Generation of 16HBEge-GFP-P2A-WT or 16HBEge-GFP-P2A-Δex23 cells

HEK293T cells were transfected with *CFTR*-containing lentiviral plasmids and packaging vector to produce retroviral particles carrying the plasmids pGFP-P2A-CFTR-WT-T7 or pGFP-P2A-CFTR-Δex23-T7. Cells were incubated for 18 hrs after transfection, then fresh medium was added to the cells. After 24 hours, the medium containing the viral particles was collected, and replaced with fresh medium. After another 24 hours, the medium containing the viral particles was collected and pooled with the previously collected medium. The pooled medium was filtered through a 0.45 μm filter. PEG-it™ solution was added to the filtered medium, with 1/5 final dilution (System Biosciences, LV810A-1), and the mixture was incubated at 4 °C for 6 hrs. The medium was centrifuged for 30 min at 1500 x g, and the supernatant was added to the 16HBE-W1282X cells. 18 hrs after the addition of viral particles, the transduced cells were selected with medium containing puromycin (0.2 μg/ml) for one week (EMD Millipore, 540411). Successful transduction was confirmed by the doxycycline-induced GFP signal in 16HBEge-GFP-P2A-WT and 16HBEge-GFP-P2A-Δex23 cells (see ‘Microscopy’).

### Plasmid transfection

All plasmids were transfected using Lipofectamine 2000 (Life Technologies, 11668019) according to the manufacturer’s protocol.

### Microscopy

#### Fluorescence microscopy

The expression of recombinant GFP-P2A-CFTR proteins in 16HBEge-GFP-P2A-WT and 16HBEge-GFP-P2A-Δex23 cells was induced with growth medium containing 0 – 2 μg/mL doxycycline for 48 hrs (Research Products International Corp, D43020-100). The images were taken with a Revolve microscope and acquired by ECHO Pro v6.0.1 software (Revolve, ECHO, San Diego, CA, United States). GFP-intensity analysis from the images was performed with ImageJ2 software (v.2.2.0).

#### Immunohistochemical microscopy

16HBE-W1282X cells were seeded on chamber slides (Corning 354118) and transfected with pGFP-P2A-CFTR-WT-T7 or pGFP-P2A-CFTR-Δex23-T7 plasmids using Lipofectamine 2000, according to the manufacturer’s protocol. 18 hrs after transfection, the culture medium was replaced with medium containing 2 μg/mL doxycycline. 48 hrs later, the cells were washed with PBS and fixed with 4% paraformaldehyde (Thermo Fisher, 28906). The fixed cells were washed with PBS and blocked with 5% goat serum for 1 hr (Thermo Fisher, 31872). Mouse anti-T7 primary antibody (Millipore Sigma, 69522) was applied at a dilution of 1:1000 for 2 hrs at room temperature. The slides were washed three times and incubated with goat anti-mouse IgG2b secondary antibody conjugated to Alexa Fluor 594 for 1 hr (Thermo Fisher, A-21145). Nuclei were stained with DAPI. Images were captured with a confocal laser-scanning microscope (Zeiss LSM710).

### ASOs transfection and treatment

All ASOs were 18mers uniformly modified with 2′-*O*-(2-methoxyethyl) (MOE) ribose, phosphorothioate linkages, and 5’-methylcytosine. They were obtained from Integrated DNA Technologies (Coralville, Iowa) or Bio-Synthesis (Lewisville, TX). The scramble control ASO is a scrambled sequence based on i23-5ss. All ASOs were dissolved in water and stored at −20 °C. Stock ASO concentrations were calculated based on the A260 and each ASO’s extinction coefficient *e* (mM^−1^ × cm^−1^ @ 260nm). The sequences of all ASOs used in this study are listed in Table 1.

**Table 1.**
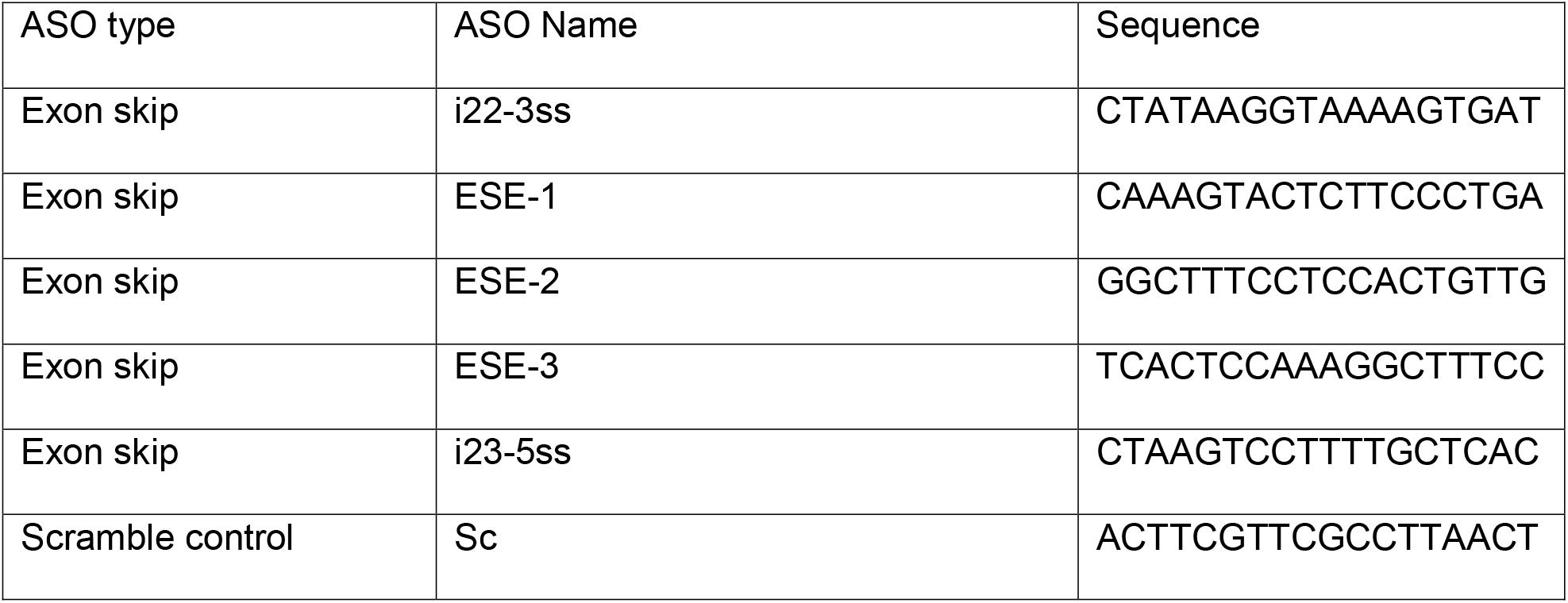
ASO sequences

Cells were transfected with ASOs and plasmids using Lipofectamine 3000 (Life Technologies, L3000015) according to the manufacturer’s protocol, and harvested 48 hrs post-transfection. For ASO treatment by free uptake, 1 mM stock ASO solutions were diluted into MEM with 10% FBS to the desired final concentrations, and the cells were cultured for 4 days before harvesting or Ussing-chamber assays. NMD-reporter expression was induced with 2 μg/mL doxycycline with culture-medium change, 6 hrs after transfection.

### RNA extraction and RT-PCR

Total RNA was extracted with TRIzol (Life Technologies) according to the manufacturer’s protocol. Oligo dT(18)-primed reverse transcription was carried out with ImProm-II Reverse Transcriptase (Roche). Semi-quantitative radioactive PCR (RT-PCR) was carried out in the presence of ^32^P-dCTP with AmpliTaq DNA polymerase (Thermo Fisher), and real-time quantitative RT-PCR (RT-qPCR) was performed with Power Sybr Green Master Mix (Thermo Fisher). Primers used for RT-PCR and RT-qPCR are listed in Table 2. RT-PCR products were separated by 6% native polyacrylamide gel electrophoresis, detected with a Typhoon FLA7000 phosphorimager, and quantitated using MultiGauge v2.3 software (Fujifilm); RT-qPCR data were quantitated using the QuantStudio 6 Flex system.

**Table 2.**
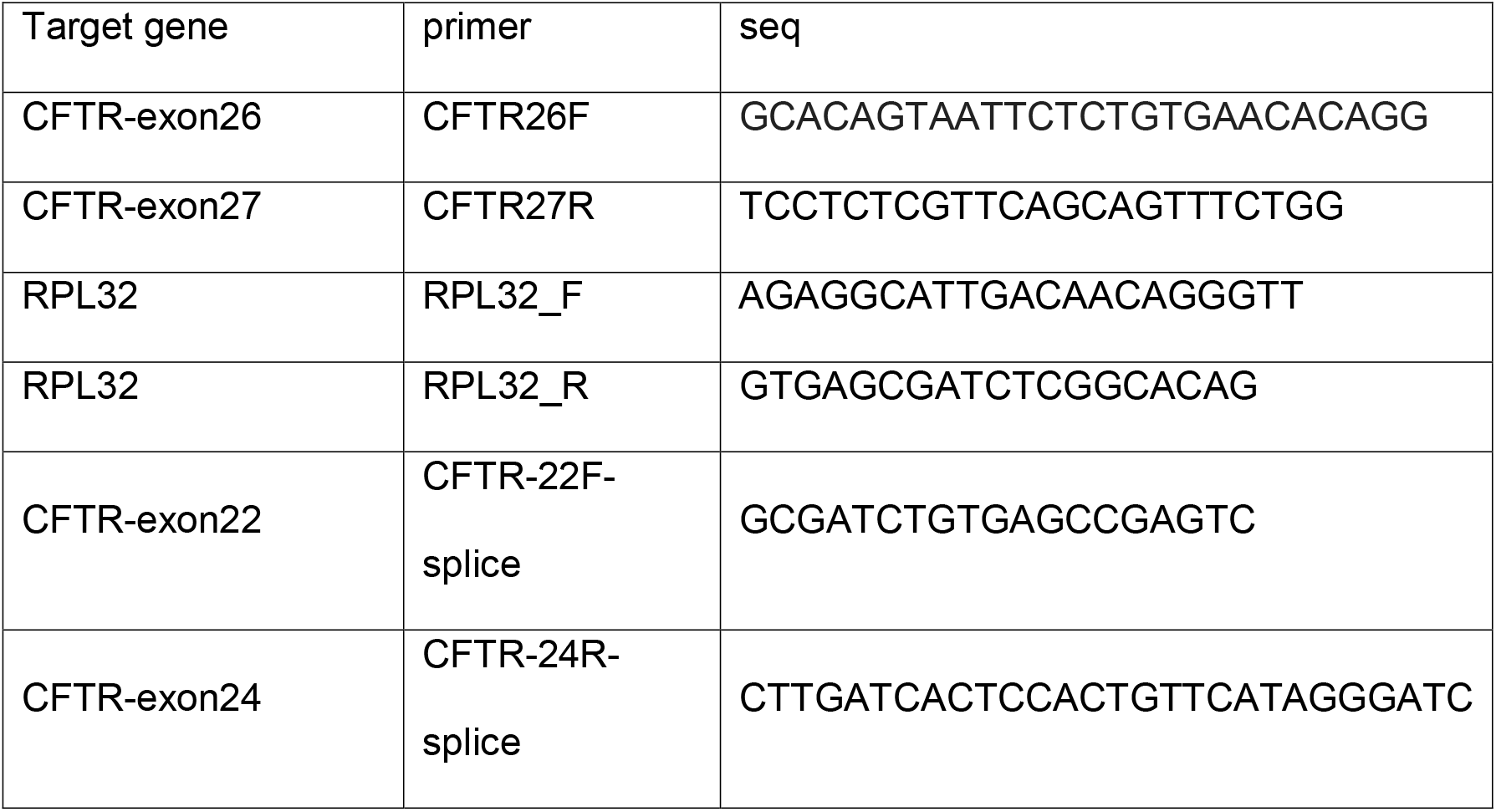
RT-PCR and qRT-PCR Primer Sequences.

### Protein extraction and Western blotting

Cells were harvested with RIPA buffer (150 mM NaCl, 50 mM Tris-HCl pH 8.0, 1% NP40, 0.5% sodium deoxycholate, and 0.1% SDS) and 2 mM EDTA + protease inhibitor cocktail (Roche) by sonicating for 5 min at medium power using a Bioruptor (Diagenode), followed by 15-min incubation on ice. Protein concentration was measured using the Bradford assay (Bio-Rad) with BSA as a standard. Cell lysates were mixed with Laemmli buffer and incubated at 37 °C for 30 min. The protein extracts were separated by sodium dodecyl sulfate-polyacrylamide gel electrophoresis (SDS-PAGE) (6% Tris-chloride gels) and then transferred onto a nitrocellulose membrane. CFTR bands C, B, and A were detected with antibody UNC-596 (J. Riordan lab, University of North Carolina, Chapel Hill, NC). The C and B band intensities were measured together for the quantification of CFTR protein levels. Na/K-ATPase, detected with a specific antibody (Santa Cruz sc-48345), was used as a loading control. IRDye 800CW or 700CW secondary antibody (LI-COR) was used for Western blotting, and the blots were imaged and quantified using an Odyssey Infrared Imaging System (LI-COR).

### Ussing-chamber assay

16HBEge cells were grown as an electrically tight monolayer on Snapwell filter supports (Corning, 3801), as described (37), and both serosal and mucosal membranes were exposed to the ASOs for 4 days, and to CFTR correctors VX-809, VX-661, or VX-445 for 24 hrs, before the assays (Selleckchem, S1565, S7059, and S8851, respectively). The Snapwell inserts were transferred to an Ussing chamber (P2302, Physiologic Instruments, Inc., San Diego, CA). The serosal side only was superfused with 5 mL of HB-PS buffer; on the mucosal side, 5 ml of CF-PS was used (137 mM Na-gluconate; 4 mM KCl; 1.8 mM CaCl_2_; 1 mM MgCl_2_; 10 mM HEPES; 10 mM glucose; pH adjusted to 7.4 with N-methyl-D-glucamine) to create a transepithelial chloride-ion gradient. After clamping the transepithelial voltage to 0 mV, the short-circuit current (I_SC_) was measured with a Physiologic Instruments VCC MC6 epithelial voltage clamp, while maintaining the buffer temperature at 37 °C. Baseline activity was recorded for 20 min before agonists (final concentrations: 10 μM forskolin (Sigma, F6886) and 10 μM VX-770 (Selleckchem, S1144) and inhibitor (final concentration: 20 μM CFTRinh-172 (Sigma, C2992)) were applied sequentially at 10-min intervals, to both serosal and mucosal surfaces. Agonists/inhibitor were added from 200x-1000x stock solutions. Data acquisition was performed with ACQUIRE & ANALYZE Revision II (Physiologic Instruments).

### ESE motif analysis

Potential SR protein binding sites were analyzed by ESEfinder and SFmap (33, 34).

### Statistical analyses

Statistical analyses were performed with GraphPad Prism 5. Statistical parameters are indicated in the figures and legends. For two-tailed t-test or one-way analysis of variance (ANOVA) with Tukey’s or Dunnett’s post-test, P<0.05 was considered significant. The Pearson correlation and P values were calculated using R. The asterisks and hash signs mark statistical significance as follows: n.s. P>0.05; *^/#^ P<0.05; **^/##^ P<0.01; ***^/###^ P<0.001.

## Acknowledgements

We are very grateful to Martin Mense and Hermann Bihler (CFFT, Lexington, MA) for generously sharing experimental protocols and advice.

## Abbreviations

ASO: Antisense oligonucleotide
CF: cystic fibrosis
CHX: cycloheximide
EJC: Exon-junction complex
ESE: Exonic splicing enhancer
GAIN: Gene-specific Antisense Inhibition of NMD
HBE: human bronchial epithelial cells
ISE: Intronic splicing enhancer
MOE: 2′-methoxyethyl
NBD: nucleotide binding domain
NMD: Nonsense-mediated mRNA decay
PS: Phosphrothioate
RBP: RNA-binding protein
RTC: Read-through compound

