## Supplementary figure for "Exon-Skipping Antisense Oligonucleotides for Cystic Fibrosis Therapy"

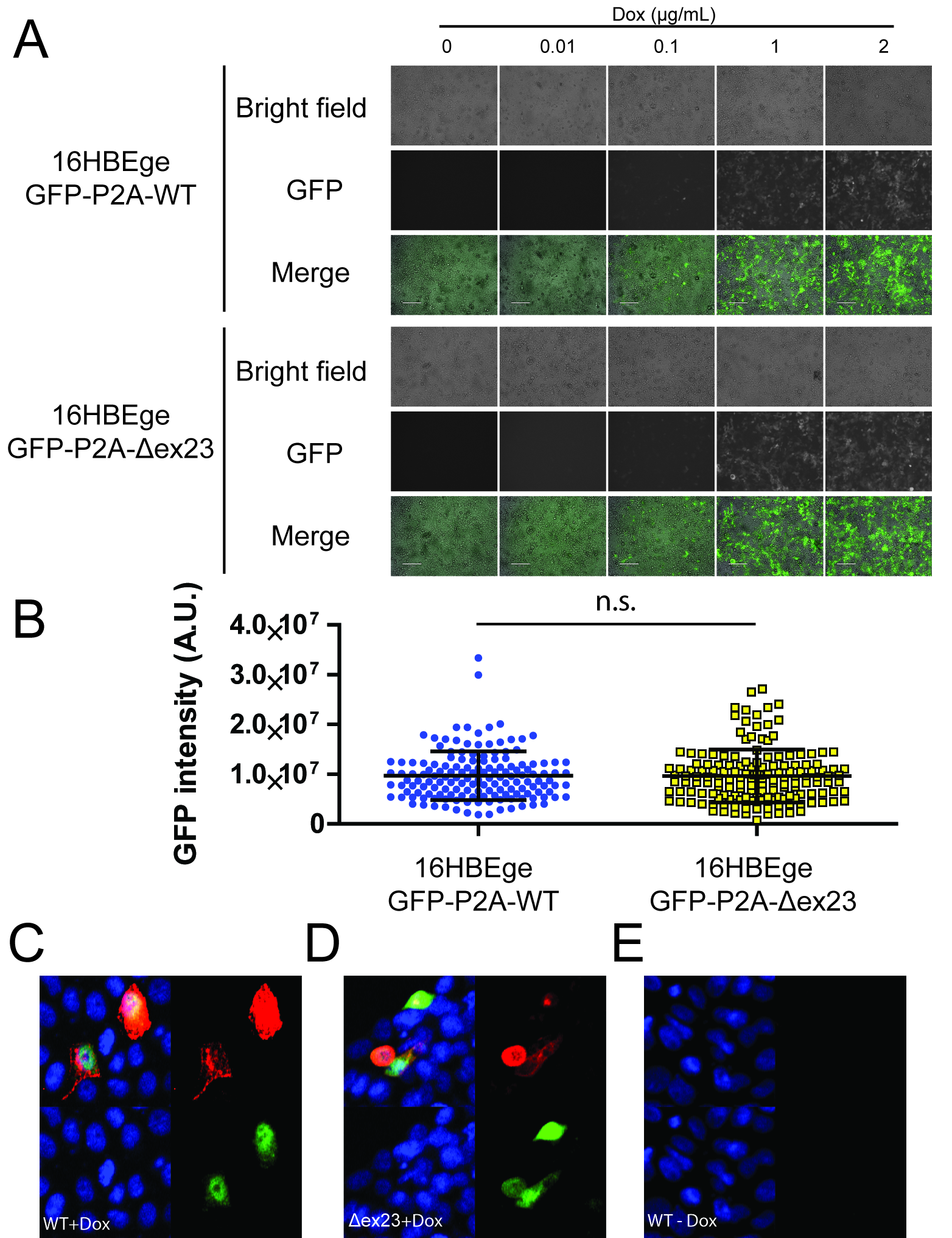

**Supplementary Figure 1. Dose-dependent induction of GFP-P2A-CFTR expression by doxycycline.**

**A.** Fluorescence microscopy images of 16HBEge-GFP-P2A-WT and 16HBEge-GFP-P2A-Δex23 cells treated with 0, 0.01, 0.1, 1, or 2 μg/mL doxycycline for 48 hrs. The white bars are 120-μm rulers. **B.** GFP intensity (arbitrary units) of individual 16HBEge-GFP-P2A-WT and 16HBEge-GFP-P2A-Δex23 cells after 48 hrs treatment with 2 μg/mL doxycycline (from panel A). (n=150 cells, Student’s t-test, n.s. P > 0.05). The longest line in the middle shows mean GFP intensity. The error bars shown above and below are standard deviations. **C-D.** (C) 16HBE-W1282X cells transfected with pGFP-P2A-WT-T7 + doxycycline. (D) 16HBE-W1282X cells transfected with pGFP-P2A-CFTR-Δex23-T7 + doxycycline. **E.** 16HBE-W1282X cells transfected with pGFP-P2A-WT-T7 without doxycycline. Upper left panel = merged image. Upper right panel = T7-epitope staining. Lower left panel = DAPI. Lower right panel = GFP.

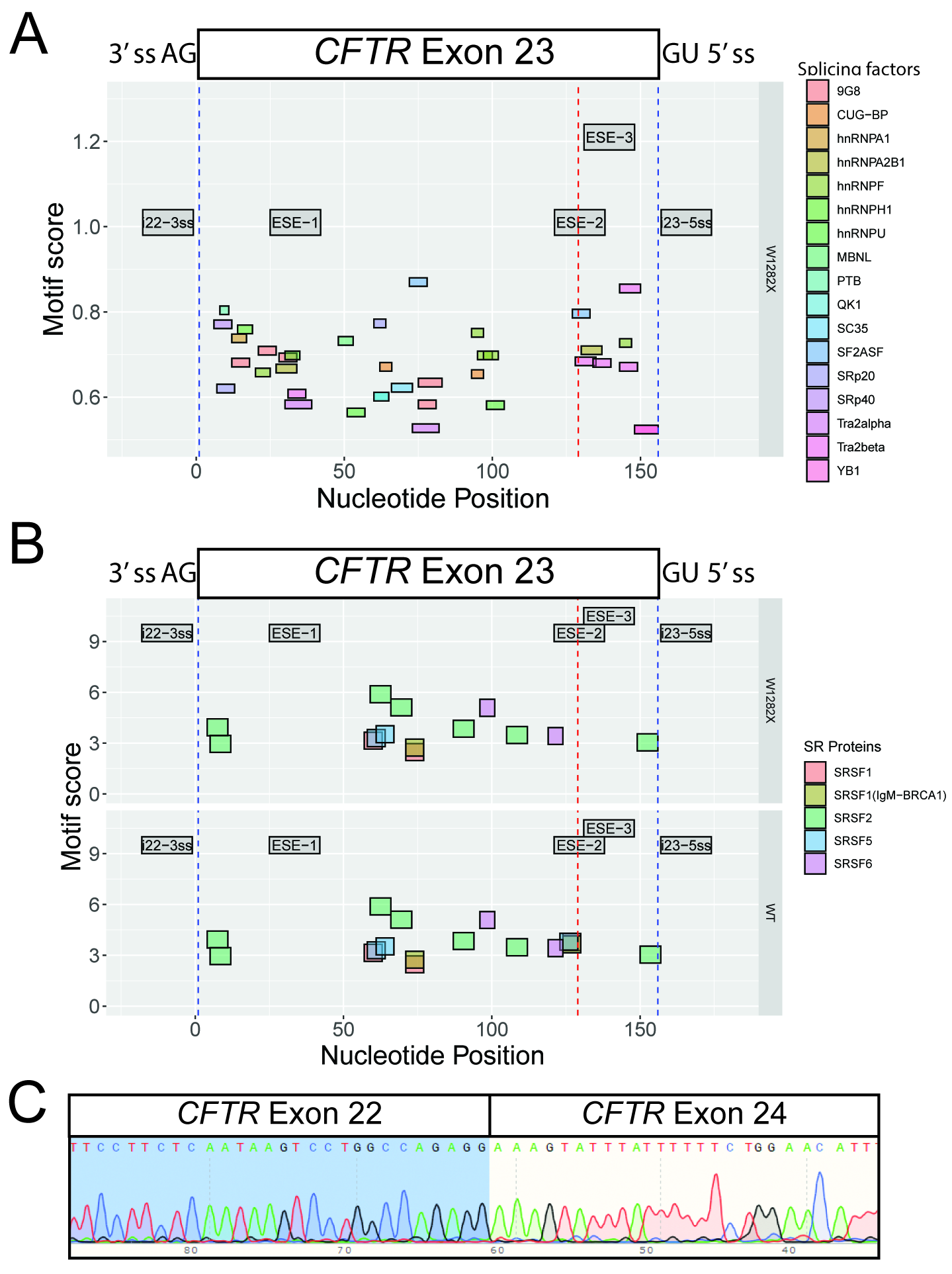

**Supplementary Figure 2. Putative ESE prediction and Sanger sequencing of Exon 23-skipped *CFTR* mRNA. A-B.** Potential splicing-factor binding sites on exon 23 were analyzed by (A) SFmap and (B) ESEfinder. The blue dashed line represents the exon 23 boundaries, and the red dashed line represents the location of the G>A mutation in W1282X. **C.** The identity of the exon-23-skipped band was confirmed by Sanger sequencing.

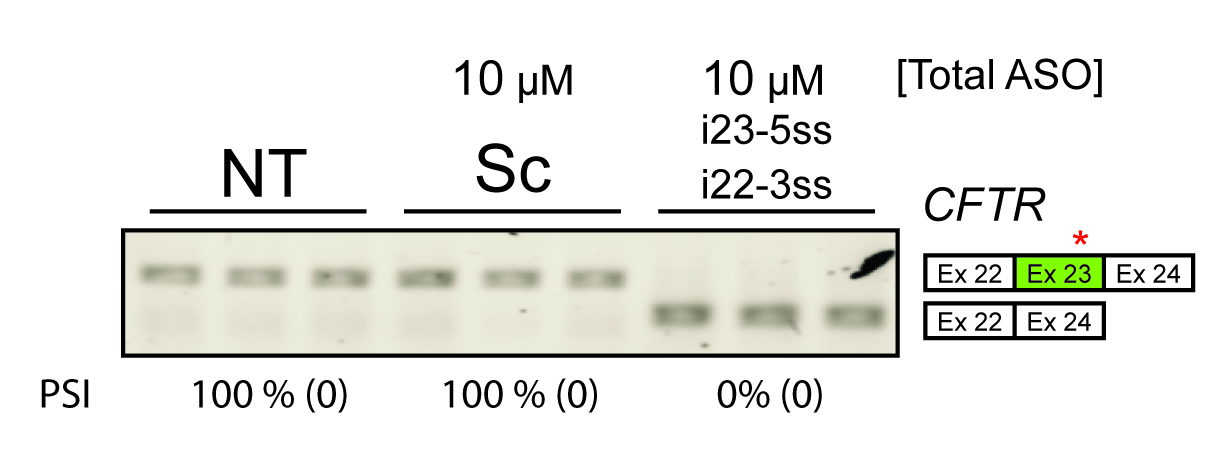

**Supplementary Figure 3. Free uptake of Exon Skipping ASO Cocktail**

Non-radioactive RT-PCR shows that free uptake treatment of 10 μM splice-site ASO cocktail promotes nearly complete exon 23 skipping in 16HBE-W1282X cells.

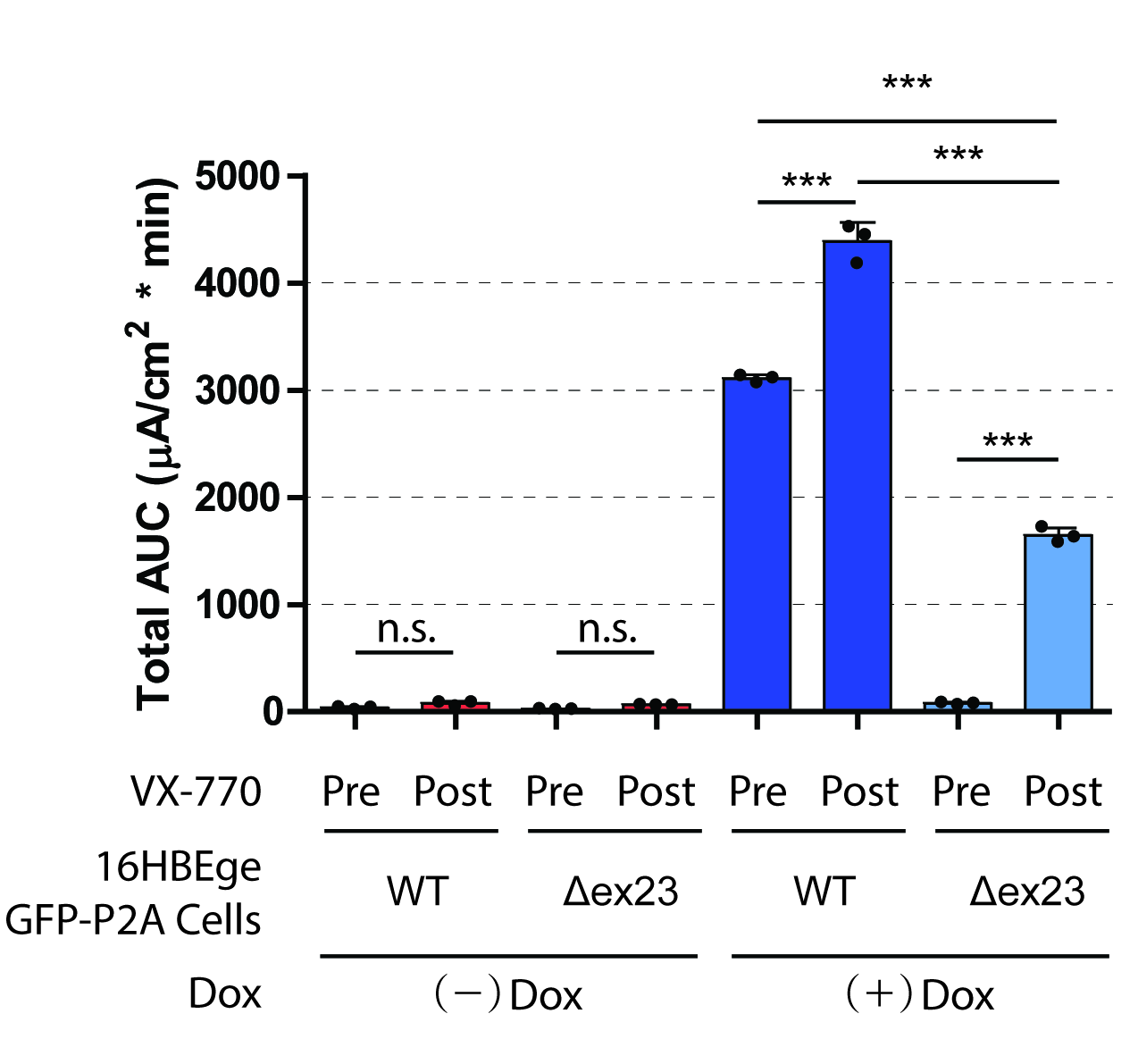

**Supplementary Figure 4. Recombinant CFTR activity in 16HBEge-GFP-P2A-WT or 16HBEge-GFP-P2A- Δex23 cells before or after VX-770 treatment.**

The total area under the curve before or after the addition of VX-770 (minute 20-30 or minute 30-40, respectively) was calculated from Fig. 1E-F (n=3 independent treatments. n.s. P>0.05, ***P<0.001, Student’s t-test). Error bars show standard deviations. Dox = doxycycline. Fsk: forskolin.
